# Leveraging information in spatial transcriptomics to predict super-resolution gene expression from histology images in tumors

**DOI:** 10.1101/2021.11.28.470212

**Authors:** Minxing Pang, Kenong Su, Mingyao Li

## Abstract

Recent developments in spatial transcriptomics (ST) technologies have enabled the profiling of transcriptome-wide gene expression while retaining the location information of measured genes within tissues. Moreover, the corresponding high-resolution hematoxylin and eosin-stained histology images are readily available for the ST tissue sections. Since histology images are easy to obtain, it is desirable to leverage information learned from ST to predict gene expression for tissue sections where only histology images are available. Here we present HisToGene, a deep learning model for gene expression prediction from histology images. To account for the spatial dependency of measured spots, HisToGene adopts Vision Transformer, a state-of-the-art method for image recognition. The well-trained HisToGene model can also predict super-resolution gene expression. Through evaluations on 32 HER2+ breast cancer samples with 9,612 spots and 785 genes, we show that HisToGene accurately predicts gene expression and outperforms ST-Net both in gene expression prediction and clustering tissue regions using the predicted expression. We further show that the predicted super-resolution gene expression also leads to higher clustering accuracy than observed gene expression. Gene expression predicted from HisToGene enables researchers to generate virtual transcriptomics data at scale and can help elucidate the molecular signatures of tissues.

## INTRODUCTION

Knowledge of the spatial organization of cells and the spatial variation of gene expression in tissues is important in understanding the complex transcriptional architecture of multicellular organisms. For example, in a heterogeneous tissue such as tumor, cancer cells can differ vastly from each other in their gene expression profiles and cellular properties due to residing in distinct tumor microenvironments. Recent advances in spatial transcriptomics (ST) technologies have made it possible to profile gene expression while retaining the spatial location information of the measured genes within tissues (1–6). Applications of the ST technologies in diverse tissues and diseases have transformed our views of transcriptome complexity (7–9). A popular ST technology is based on spatial barcoding followed by next-generation sequencing in which transcriptome-wide gene expression is measured in gene capture locations, referred to as spatially barcoded spots. Such ST technologies include Spatial Transcriptomics (10), 10x Genomics Visium, SLIDE-seq (11), SLIDE-seq2 (12), and many others (13,14). Data from such spatial barcoding-based technology typically include a high-resolution hematoxylin and eosin (H&E)-stained histology image of the tissue section from which the gene expression data are obtained.

Although ST offers rich information, such data are still expensive to generate, which prevents the applications of ST in large-scale studies. On the other hand, H&E-stained histology images are easier and cheaper to obtain than ST and are routinely generated in clinics. It is desirable to leverage information learned from ST to predict gene expression from histology images. Such predictions can generate virtual ST data, which will enable the study of spatial variations of gene expression at scale. Indeed, several studies have shown that tumor related genes are highly correlated with histological features, suggesting that gene expression can be predicted from histology images. HE2RNA (15), a model based on the integration of multiple data modes, is trained to systematically predict gene expression profiles from whole-slide images without the reliance on expert annotation. ST-Net (16) predicts spatially variable gene expression from histology images using a supervised convolutional neural network (CNN) trained from breast cancer ST data.

While these methods have shown promising performance, they are not short of limitations. HE2RNA was developed for bulk RNA sequencing and lacks the ability to learn from ST data. While ST-Net is specifically designed for ST, it does not model the spatial location information of each measured spot in their CNN model. Since gene expression often displays local patterns, which are often manifested in the histology images, it is necessary to explicitly model the spatial location information when predicting gene expression. Although CNN-based models have dominated the field of computer vision for years, different parts of an image must be processed in the same way regardless of their position. This intrinsic bias in CNN makes it less ideal for ST data. However, this bias has been recently alleviated by Vision Transformer (17), which internally utilizes self-attention mechanism for divided image patches and has shown strong performance on many tasks, including medical image classification, segmentation (18), and registration (19).

To utilize these advances in Vision Transformer, we developed HisToGene, an attention-based model that aims to predict gene expression from H&E-stained histology images based on the relationship between histological features and gene expression features learned from a training ST dataset. To account for the spatial dependency of measured spots in ST, HisToGene employs a modified Vision Transformer model, which can naturally model the positional relationship between spots through appropriate positional embedding. Compared to ST-Net (16), our attention-based model considers the spot dependency together with histological features when predicting gene expression. After model training, HisToGene can further predict super-resolution gene expression by averaging predicted gene expression from densely sampled histology image patches. To the best of our knowledge, it is the first time that gene expression can be predicted at such high resolution based on histology images. Gene expression predicted from HisToGene enables researchers to generate virtual transcriptomics data at scale and can help elucidate the molecular signatures of tissues.

## MATERIALS AND METHODS

### Overview of HisToGene

HisToGene takes a ST dataset, possibly with multiple tissue sections obtained from different subjects, as input for model training. For each tissue section in the ST data, it starts by extracting patches from the histology image according to the spatial coordinates and size of each spot in the ST data. The patch embedding and position embedding are then aggregated by learnable linear layers through a modified Vision Transformer model. Next, HisToGene utilizes multi-head attention layers to generate latent embeddings (**Figure. 1a**). With the well-trained model, HisToGene can predict gene expression for each sampled patch from the histology image in a test dataset that only has histology images. Furthermore, using a dense image patch sampling strategy, HisToGene can predict super-resolution gene expression with 4 times of the original patch/spot level resolution by default (**Figure. 1b**), but the resolution can be increased to an even higher level when using more densely sampled patches.

**Figure 1.**
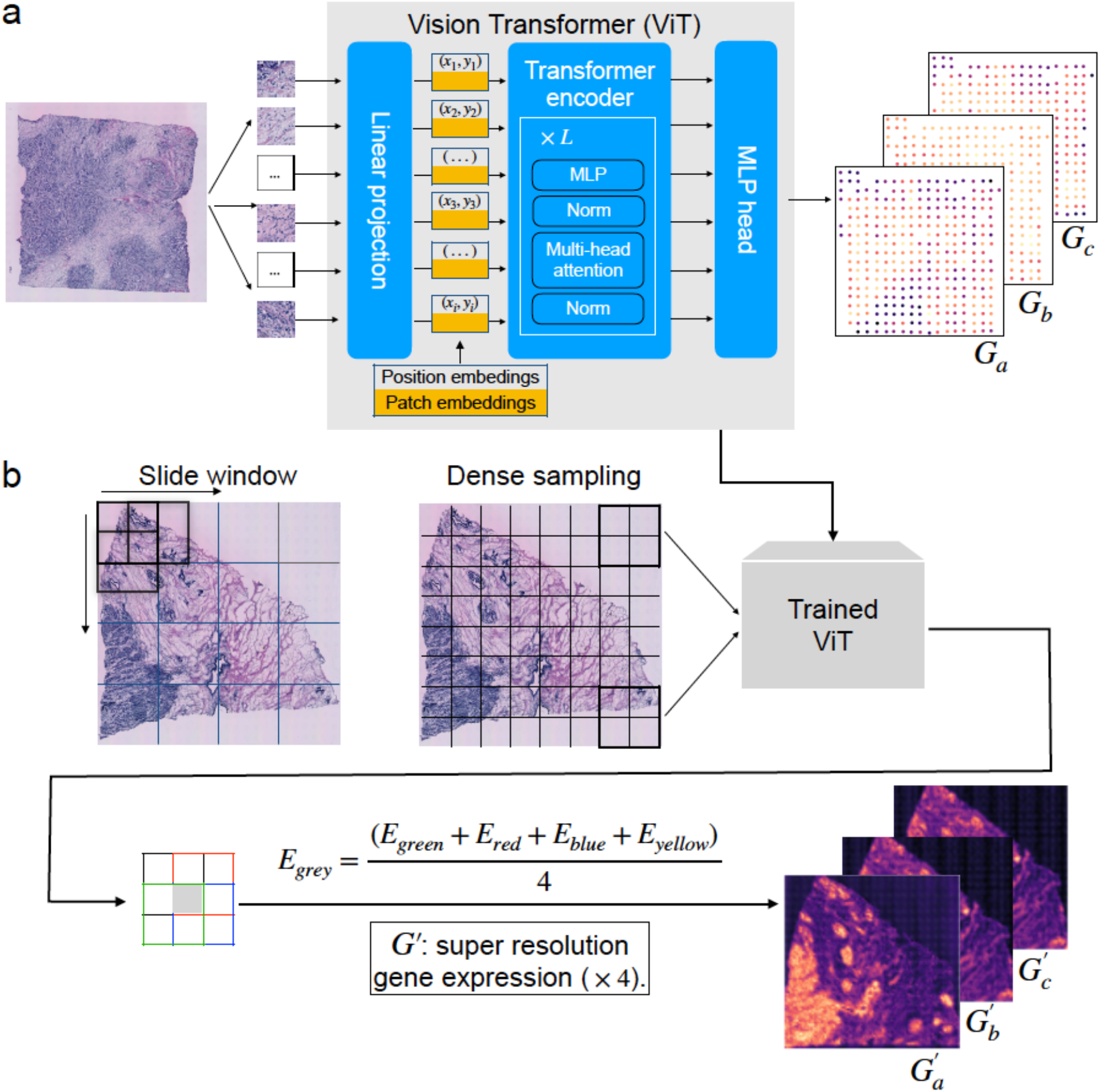
Workflow of HisToGene. **a,** The modified Vision Transformer in HisToGene starts from sequences of extracted patches from histology images in the training ST dataset. Added by the position embedding obtained from the spatial coordinates of the spots, the patch embedding goes through the Multi-Head Attention layers and is transformed by a linear layer. The final output of the modified Vision Transformer is the spatial gene prediction. This modified Vision Transformer will be trained based on the observed gene expression in the training ST dataset. **b,** Using the trained model, HisToGene can predict super-resolution gene expression. HisToGene first predicts gene expression for each sampled patch from the histology image in a test dataset that only has histology images. Using a dense image patch sampling strategy, HisToGene then predicts the super-resolution gene expression by averaging the predicted gene expression among overlapping patches. With the patch sampling strategy shown in **b**, the resolution is increased 4 times, but the resolution can be increased to an even higher level when using more densely sampled patches.

### Data preprocessing

HisToGene involves a training step and a prediction step. The training step takes a ST dataset as input, which includes histology images, the gene expression data, and the spatial coordinates for the spatial barcodes. In the training stage, it uses the histology images and the spatial coordinates of the spatially barcoded spots as input, and the corresponding spatial gene expression data as labels. The spatial gene expression data are stored in an *N* × *D* matrix of unique molecular identifier (UMI) counts with *N* spots and *D* genes. The coordinates of the spots are stored in an *N* × 2 matrix indicating the (*x, y*) location of each spot.

For the histology image, we extract patches according to the size and location of each spot. We assemble all patches in a tissue section and flatten them into an *N* × (3 × *W* × *H*) matrix as the input for the Vision Transformer, where 3 is the number of channels, and *W* and *H* represent the width and height of the patch. In our experiment on the HER2+ breast cancer dataset (20), *W* = *H* = 112 pixels, which correspond to the diameter of each spot in the ST data.

For the spatial gene expression data, we first identify common genes across all tissue sections in the training ST data. We then select the top 1,000 highly variable genes in each tissue section and eliminate genes that are expressed in less than 1,000 spots across all tissue sections. The gene expression values in each spot are normalized such that the UMI count for each gene is divided by the total UMI counts across all genes in that spot, multiplied by 1,000,000, and then transformed to a natural log scale.

### The modified Vision Transformer model for gene expression prediction

#### Model architecture

Vision Transformer has been widely used in computer vision for image recognition and outperformed other state-of-the-art methods in the ImageNet Large-Scale Visual Recognition Challenge. The standard Vision Transformer model splits an image into a fixed number of patches. However, in ST data, the number of spots that cover the captured tissue area is not fixed. This property is similar to problems in natural language processing in which the lengths of sentences are also variant. To accommodate variable numbers of spots in ST, we redesign the encoding part of the Vision Transformer model with details described below.

#### Encoding of histology image and position features

After preprocessing, we use a learnable linear layer ***W**_h_* to transform the histology image features from an *N* × (3 × *W* × *H*) matrix ***F**_h_* to an *N* × 1024 matrix ***E**_h_*, i.e., ***E**_h_* = ***F**_h_* · ***W**_h_*. Another part of the input is the *N* × 2 matrix, which represents the (*x, y*) coordinates of each spot in the ST data. The *x*-coordinate information is represented by an *N* × 1 vector, which is transformed into a one-hot encoding matrix ***P**_x_* with size *N* × *m*, where *m* is the maximum number of *x*-coordinates among all tissue sections. For the HER2+ breast cancer dataset, *m* = 30. Next, we use a learnable linear layer ***W**_x_* ∈ R^30×1024^ to transform ***P**_x_* into an *N* × 1024 matrix ***E**_x_*, i.e., ***E**_x_* = ***P**_x_* · ***W**_x_*. Similar transformations are performed for the *y*-coordinate vector to obtain an *N* × 1024 encoding matrix ***E**_y_*. Finally, we obtain the embedding matrix by summing up the image feature encoding matrix, the *x*-coordinate encoding matrix, and the *y*-coordinate encoding matrix,

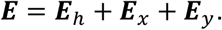

This embedding matrix is used as the input for the multi-head attention layers as described below.

#### Multi-Head Attention layers

The Multi-Head Attention module can automatically learn the attention for a “sequence”. In language data, the “sequence” is sequence of words in a sentence. In ST data, the “sequence” is a sequence of spots/patches in a tissue section. The multi-head attention is a linear combination of multiple attention heads,

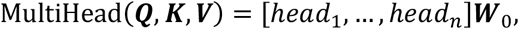

where ***W***_0_ is a learnable 1024 × 1024 parameter matrix that is used to aggregate the attention heads, *n* is the number of heads, and ***Q, K, V*** represent Query, Key, and Value. In our model, the input matrix is the *N* × 1024 embedding matrix ***E*** obtained in the previous step. The attention mechanism is defined as

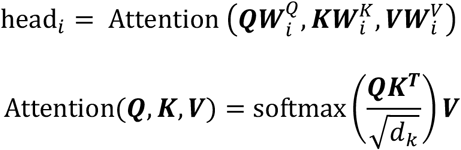

where 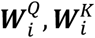, and 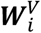 are all learnable 1024 × 1024 parameter matrices. The shape of the input for the attention is *N* × 1024. In the attention operation, we have two parts, 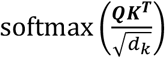 and ***V***. The former part is called Attention Map, whose shape is *N* × *N*. The latter part is the Value of the self-attention mechanism, where ***Q*** = ***K*** = ***V***. Each column of the Attention Map represents the attention weight contributed from other spots. The Attention Map provides useful information on how the model works. The result of the attention operation is an *N* × 1024 matrix. We use the output of each multi-head Attention layer as the input for the next layer and repeat this calculation sequentially.

#### Details of the model implementation

We implement the HisToGene model using PyTorch with the following hyper-parameters: learning rate is 10^-5^, the number of training epochs is 100, drop-out ratio is 0.1, the number of Multi-Head Attention layers is 8, and the number of attention heads is 16.

### Predicting gene expression at super-resolution

The above trained Vision Transformer model can predict gene expression from histology images with spot level resolution as the training ST data only contain gene expression measured within spatially barcoded spots. However, since the histology image does not have tissue gaps, it is possible to densely sample histology image patches and use predicted gene expression from overlapping patches to estimate gene expression at a resolution that is higher than the original spot. This is analogous to natural language processing, where the Transformer is trained using short sentences but can make predictions for long sentences. In our case, the “sentence” is the sequence of “spots”. Therefore, using the trained model, we can predict the expression for more “spots” within a tissue section.

The key for our super-resolution gene expression prediction lies in the dense sampling scheme of the histology image patches. First, we uniformly sample patches from the tissue area according to a grid with size of each cell determined by the spot size in the training ST data. For example, in the HER2+ breast cancer dataset, each patch is 112×112 pixels. We sample patches from the histology image such that the entire tissue area is covered by non-overlapping patches of size 112×112 pixels. By sampling the patches this way, each sub-patch is covered by 4 patches. Using the trained model, we can predict the gene expression for each sampled patch. For each sub-patch, its gene expression is predicted by the average of the predicted expressions for the 4 patches that cover it. Since the size of each sub-patch is only ¼ of the original patch, this sampling scheme can increase the gene expression resolution by 4 times. We note that our sampling scheme can be easily modified to increase gene expression prediction resolution to a higher level.

## RESULTS

### Overview of evaluations

To evaluate the performance of HisToGene, we analyzed the HER2+ breast cancer dataset (20), which includes 36 tissue sections collected from 8 HER2+ breast cancer patients. We retained 32 sections from 7 patients that have at least 180 spots per section in the analysis. To evaluate the gene expression prediction accuracy, we conducted leave-one-out (32-fold) cross validation. Specifically, for each section, we used the other 31 sections to train the model and make spatial gene expression predictions for that section. To select genes for prediction, we first considered the top 1,000 highly variable genes for each section and then filtered those that were expressed in less than 1,000 spots across all tissue sections. This filtering left with 9,612 spots and 785 genes for model training. We compared HisToGene with ST-Net for gene expression prediction. Since the source codes of ST-Net released by the authors are not maintained, we were not able to analyze the HER2+ breast cancer data using their codes. For comparison, we used our own implementation of ST-Net.

### HisToGene enables super-resolution gene expression prediction and consistently outperforms ST-Net

Since there are no tissue gaps in a histology image, it is possible to densely sample patches from the image, predict gene expression for each sampled patch, and then use the average of the predicted expression from overlapping patches to predict the gene expression for the overlapping tissue area. This allows us to increase the gene expression prediction resolution as the overlapping area among patches is much smaller than the size of the original patch. By averaging predicted gene expression across spatially close patches also reduces prediction uncertainty. Based upon this intuition, we implemented a super-resolution gene expression prediction algorithm in which the modified Vision Transformer in HisToGene can take image patches with variable lengths as input. With the patterned dense sampling of image patches shown in **Figure 1b**, we can increase the gene expression prediction resolution by 4 times. Using a similar patterned image patch sampling scheme, the gene expression prediction resolution can be increased by 9 times, 25 times, or higher.

For illustration, we sampled in the image patches such that the gene expression resolution prediction is increased by 4 times. An ideal super-resolution gene expression prediction method should increase the gene expression resolution while retaining the original expression pattern at the patch level, i.e, spot level, as this will ensure no artificial patterns are introduced during the super-resolution gene expression prediction. To evaluate whether HisToGene has this property, we obtained the patch/spot level gene expression from the super-resolution expression predicted by HisToGene. Specifically, we summed up the expression values for 4 adjacent sub-patches to “recover” the patch/spot-level gene expression. Results obtained from this super-resolution expression “recovered” approach were denoted by HisToGene*. We conducted the leave-one-out cross validation for the 32 tissue sections in the HER2+ breast cancer dataset. For each tissue section, we calculated the correlations between the observed gene expression and the predicted gene expression. **Figure 2a** shows that among the 32 tissue sections, HisToGene* predicted patch/spot level gene expression has significantly higher correlations with the observed spot-level gene expression than HisToGene for 19 (59%) sections, whereas HisToGene has significantly higher correlations than HisToGene* for 6 (19%) sections. These results indicate that with the densely sampled image patches as input in the trained prediction model, we can not only increase gene expression prediction resolution, but also the patch/spot-level gene expression prediction accuracy. Such increased accuracy is due to the flexibility of the attention mechanism in handling longer sequences of image patches, which makes the prediction benefit from information in additional batches included in the longer sequences. The increased accuracy is also due to the use of average predicted expression across nearby patches as the random error of the mean is less than that of an individual prediction.

**Figure 2.**
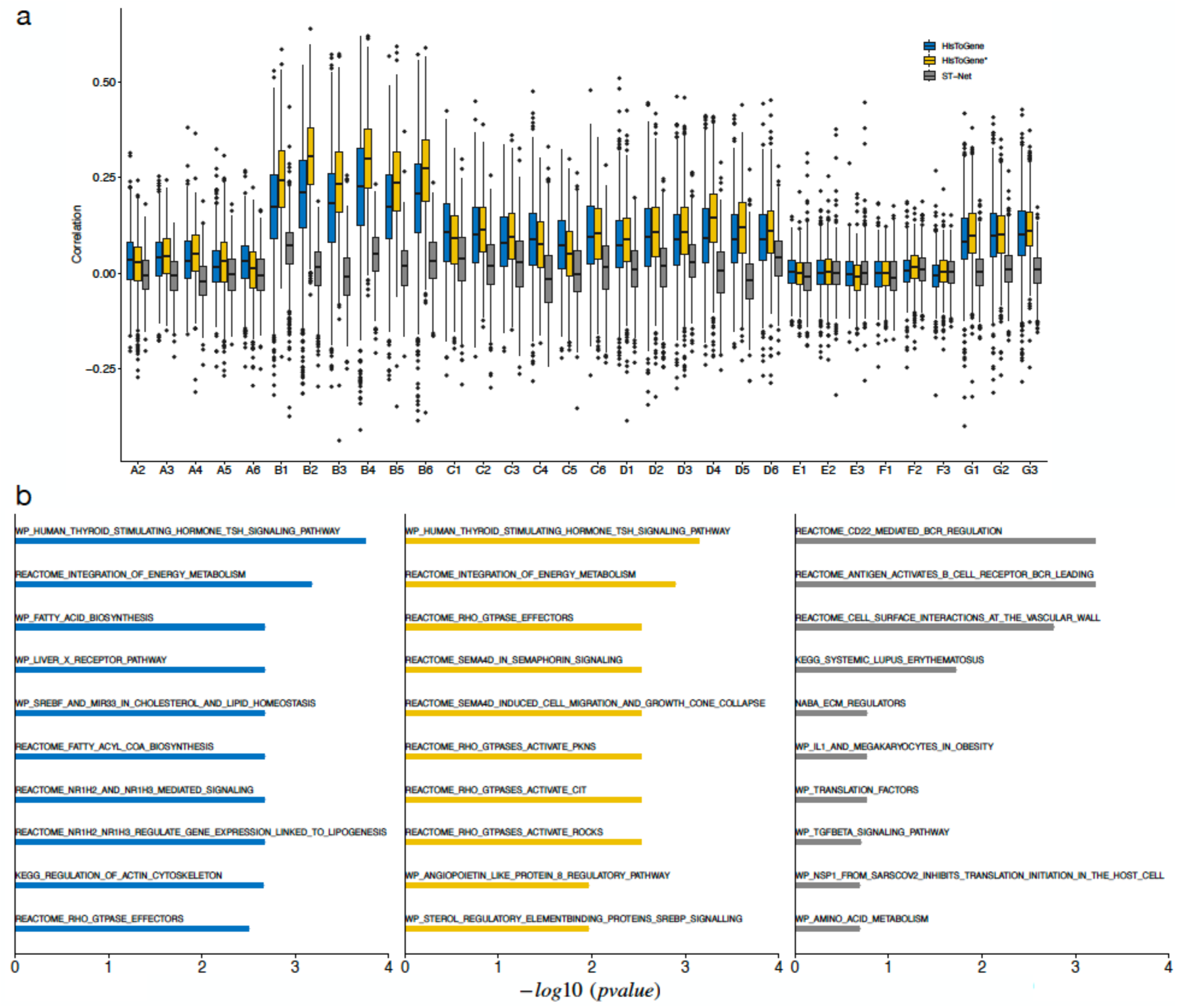
Evaluation of gene expression prediction for the HER2+ breast cancer dataset. **a,** Boxplot of the Pearson correlations between the predicted and observed gene expression for the 785 genes predicted by HisToGene, HisToGene*, and ST-Net. HisToGene* is based on the recovered patch/spot level gene expression obtained from the super-resolution gene expression prediction, denoted by HisToGene_SR. **b,** Enrichment analysis for the top 100 predicted genes by HisToGene, HisToGene*, and ST-Net.

We also performed gene expression prediction using ST-Net but found its predictions generally yielded low correlations with the observed expression. In fact, for most of the tissue sections, the mean correlations are around zero, and the correlations are not much better even for patient B in which both HisToGene and HisToGene* yielded much higher correlations. We suspect the relatively poor performance of ST-Net is due to its failure in considering the spatial dependency of spots when building the prediction model. As such, patches obtained from different patients are treated in the same way. As reported in the original study (20), there are strong subject-to-subject differences among patients, thus ignoring such differences would lead to less accurate prediction. By contrast, the modified Vision Transformer in HisToGene considers a tissue section as the modeling unit, thus the histology and gene expression relationships are learned within each tissue section, which helps alleviate the subject-to-subject differences among patients. These results demonstrate the importance of considering the spatial dependency of spots when training the prediction model.

To show that both HisToGene and HisToGene* can predict biologically meaningful information, we conducted gene set enrichment analysis using fgsea (21). Inspired by iPath (22), which sorts genes by positive values, for each approach, we ranked the genes by the average -log10 p-values across all 32 tissue sections, where the p-value for each tissue section was obtained by testing whether the correlation between the observed and the predicted expression values was significantly different from zero. We used the top 100 genes to calculate the enrichment score for each pathway from the C2 canonical pathways in MSigDB (23). Then, the significance for each pathway was assessed by permutations (n=10,000) of the gene list. The enrichment analysis results demonstrate that the highly correlated genes in HisToGene and HisToGene* are enriched in breast-cancer-related pathways (**Figure 2b**). For example, HisToGene*’s top enriched pathways include human thyroid stimulating hormone pathway and REACTOME integration of energy metabolism pathway. Previous studies have reported that thyroid hormones are associated with the risk of breast cancer (24), and energy metabolism is a hallmark of cancer cells and links with the breast cancer brain metastases (25). By contrast, the top enriched pathways for ST-Net show less relevance with breast cancer.

### Visualization of the predicted gene expression

To gain a better understanding of the predicted gene expression, we next selected the top predicted genes obtained from each method for visualization. For each gene in a tissue section, we calculated the correlation between the observed and the predicted expression values and tested whether the correlation is significantly different from zero. We then ranked the genes by the average -log10 p-values across all 32 tissue sections. **Figure 3a** (**Supplementary Table 1**) shows the top 4 genes (*GNAS, MYL12B, FASN, and CLDN4*) predicted by HisToGene, where the expression for the best predicted tissue section by HisToGene was visualized. *GNAS* (mean R = 0.32) encodes the stimulatory G-protein alpha subunit and regulates the production of the second messenger cyclic AMP. Elevated expression of *GNAS* has been found in several tumors including breast cancer and promotes breast cancer cell proliferation (26). *MYL12B* (mean R = 0.27) encodes a myosin regulatory subunit that plays an important role in the regulation of non-muscle cell contractile activity via its phosphorylation. A recent study showed that the activity of myosin II in cancer cells drives tumor progression, where the activation of myosin II in non-muscle cells is regulated by phosphorylation of a regulatory light chain such as *MYL12B* (27). *FASN* (mean R = 0.27) encodes a key enzyme that is involved in the biogenesis of membrane lipids in proliferating cells and is closely associated with the occurrence and development of tumors (28). Inhibition of *FASN* induces apoptosis in breast cancer cells, making it a potential therapeutic target for breast cancer (29). *CLDN4* (mean R = 0.26) encodes a tight junction protein that is required for cell adhesion. It is frequently expressed in primary breast cancers, especially in their metastases, thus is a promising membrane bound molecular imaging and drug target for breast cancer (30–32). As a comparison, we also included the predicted gene expression obtained from HisToGene*, the super-resolution gene expression (denoted by HisToGene_SR), and ST-Net. Although the mean -log10 p-values for genes obtained from HisToGene* are not as significant as HisToGene, the general predicted expression patterns are similar to HisToGene, and for the selected tissue sections, the correlations are similar in magnitude to HisToGene. By contrast, the ST-Net predicted expression shows little correlation with the observed expression.

**Figure 3.**
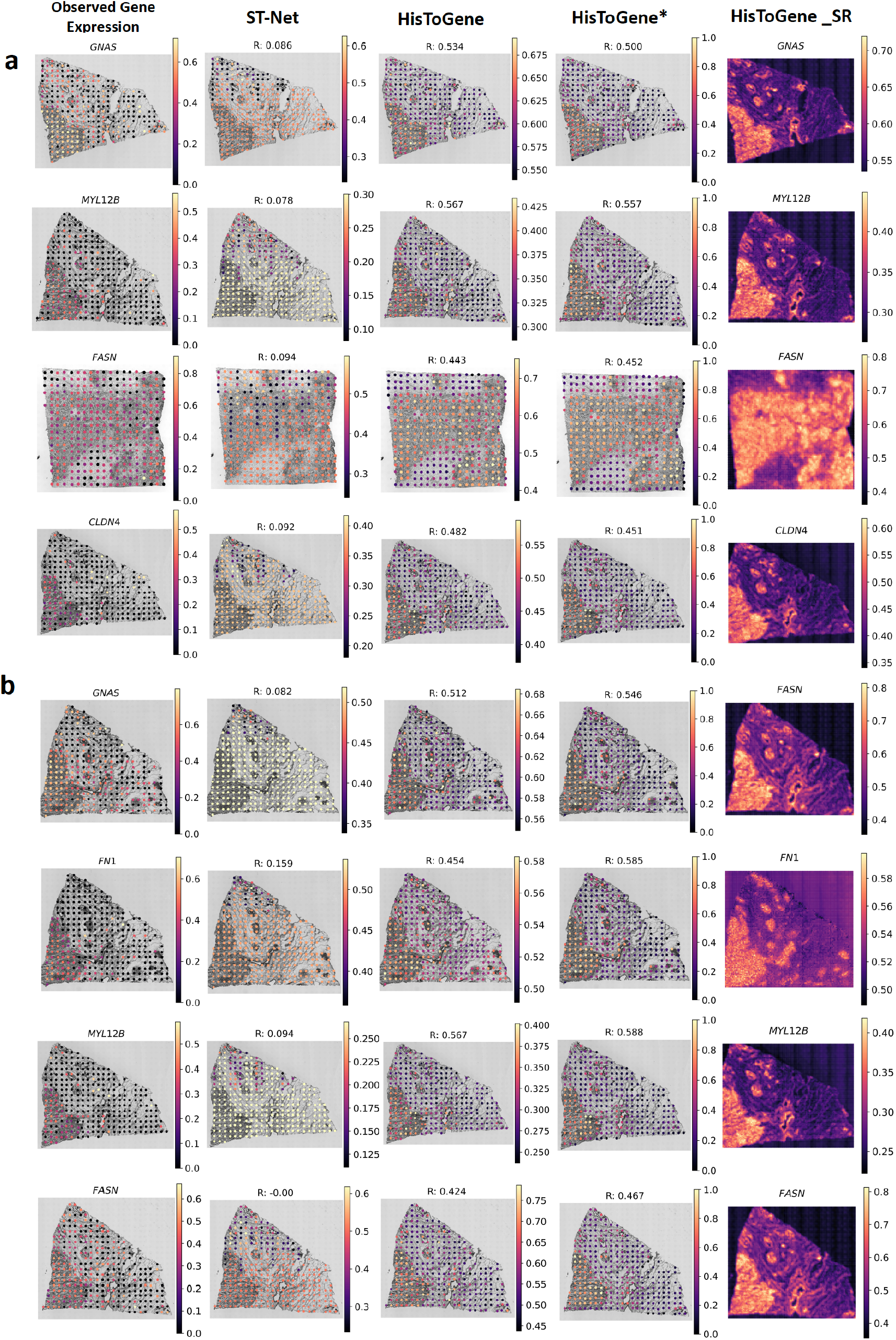
Visualization of the top predicted genes in the HER2+ breast cancer dataset. **a,** Relative expression of the top 4 genes predicted by HisToGene. The genes were selected based on the average -log10 p-values across all 32 tissue sections, where the p-value for each tissue section was obtained by testing whether the correlation between the predicted and observed gene expression was significantly different from zero. For each of the 4 genes, the tissue section that had the smallest p-value by HisToGene was selected for visualization. HisToGene* was based on the recovered patch/spot level gene expression obtained from the super-resolution gene expression prediction, denoted by HisToGene_SR. **b,** Relative expression of the top 4 genes predicted by HisToGene*. The genes were selected based on the average -log10 p-values across all 32 tissue sections, where the p-value for each tissue section was obtained by testing whether the correlation between the predicted and observed gene expression was significantly different from zero. For each of the 4 genes, the tissue section that had the smallest p-value by HisToGene* was selected for visualization. HisToGene* was based on the recovered patch/spot level gene expression obtained from the super-resolution gene expression prediction, denoted by HisToGene_SR.

**Figure 3b (Supplementary Table 2)** shows the top 4 genes (*GNAS*, *FN1*, *MYL12B*, and *FASN*) predicted by HisToGene*, 3 of them (*GNAS*, *MYL12B*, and *FASN*) were also predicted by HisToGene as the top genes. *FN1* is a gene that shows higher correlation in HisToGene* (mean R = 0.24) than in HisToGene (mean R = 0.22). *FN1* encodes fibronectin, a glycoprotein that is present in a soluble dimeric form in plasma, and in a dimeric or multimeric form at the cell surface and in extracellular matrix. Fibronectin is involved in cell adhesion and migration processes, and high expression of *FN1* is associated with breast cancer invasion and metastasis (33). Interestingly, although *FASN* is among the top 4 best predicted genes by both HisToGene* (mean R = 0.24) and HisToGene (mean R = 0.27), the best predicted tissue sections are different. For the best tissue section predicted by HisToGene* (R = 0.47), the HisToGene correlation is 0.42, only slightly worse than HisToGene*, whereas the correlation obtained from ST-Net prediction is close to 0. In general, we found that HisToGene* has higher correlations than HisToGene, whereas the correlations for ST-Net are often close to 0. For *GNAS, FN1, MYL12B*, and *FASN*, we further examined the super-resolution gene expression prediction, which revealed fine grained spatial expression patterns that are missed in the original patch/spot level gene expression prediction.

As a comparison, we also visualized the top 4 genes (*IGHM, PPP1R1B, IGLC2*, and *PNMT*) predicted by ST-Net (**Supplementary Figure 1 and Supplementary Table 3**). The average correlations for these 4 genes are much lower than the top 4 genes predicted by HisToGene and HisToGene*.

### HisToGene predicted gene expression can recover pathologists annotated spatial domains

Next, we examined if the predicted gene expression can be used to recover the pathologists annotated spatial domains. The HER2+ breast cancer data included 6 tissue sections with pathologists’ annotation, which allowed us to further evaluate if the predicted gene expression patterns are biologically meaningful. If the predictions are useful in revealing the underlying biology, we would expect the clusters obtained using the predicted gene expression to agree well with the pathologists annotated spatial domains. We performed K-Means clustering using the predicted gene expression obtained from HisToGene, HisToGene*, and ST-Net. The clustering results were evaluated using Adjusted Rand Index (ARI) by treating pathologists annotated spatial domains as the ground truth. As a comparison, we also performed clustering analysis using the observed gene expression for each tissue section.

**Figure 4** shows the clustering results and the corresponding ARIs for each method and the results obtained using the observed gene expression. Among the 6 tissue sections with pathologists’ annotation, HisToGene* yielded the highest ARIs for 4 sections (B1, C1, D1, and F1), and for sections D1 and F1, the HisToGene*’s ARIs are much higher than those obtained from the observed gene expression and ST-Net. For E1, ST-Net had the highest ARI. For G2, the observed gene expression had the highest ARI. Clustering analysis using observed gene expression is a commonly conducted task in spatial transcriptomics (34–36). Interestingly, HisToGene* had even higher ARIs than the observed gene expression for 4 out of the 6 tissue sections. Since HisToGene* is based on the aggregated super-resolution gene expression, we next performed clustering analysis using the super-resolution gene expression, denoted by HisToGene_SR. Although we cannot directly calculate the ARIs for HisToGene_SR, visual examination indicates that the clustering results agreed well with the pathologists annotated spatial domains, with the tumor region clearly separated from the background.

**Figure 4.**
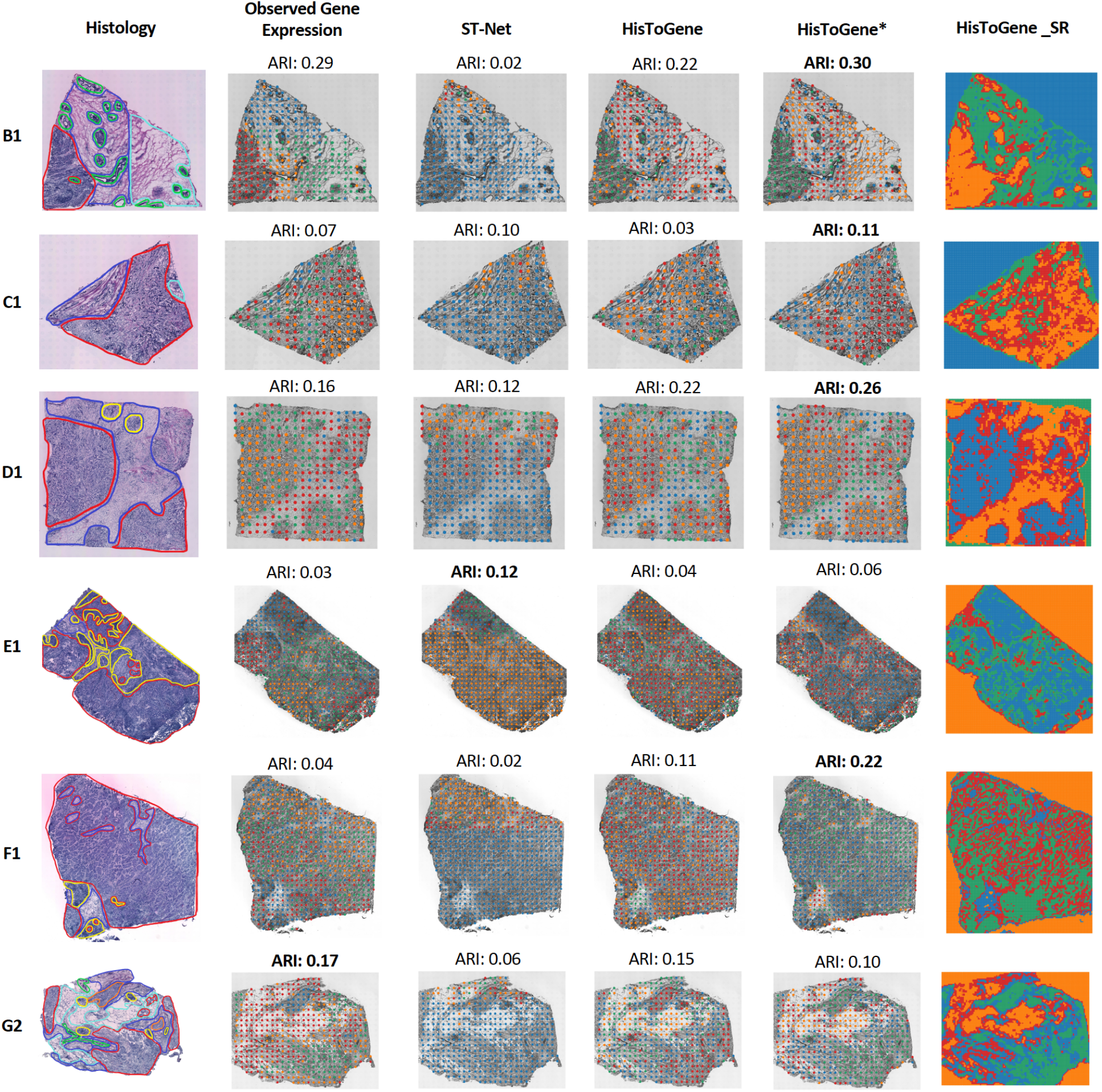
Clustering analysis using predicted gene expression in the HER2+ breast cancer dataset. 6 of the 32 tissue sections had pathologists’ annotation which allowed us to evaluate whether the predicted gene expression can reveal the pathologists annotated spatial domains. The first column shows the histology image with the pathologists’ annotation, where the red lines represent invasive cancer, green lines represent breast glands, yellow lines represent immune infiltrate, and the blue lines represent connective tissue. The remaining columns show the clustering results generated from the observed, ST-Net predicted gene expression, HisToGene predicted gene expression, HisToGene* predicted gene expression, and HisToGene_SR using K-Means clustering algorithm (k=4). Clustering accuracy was evaluated by the Adjusted Rand Index (ARI) between the pathology annotations and the clusters obtained from the predicted gene expression.

### Understanding the HisToGene prediction with attention map

It is intriguing that the predicted super-resolution gene expression led to higher clustering ARIs than the observed gene expression. We next sought to investigate how the super-resolution gene expression prediction works. Attention is a key feature in HisToGene’s modified Vision Transformer model. To understand how attention contributes to the HisToGene predicted gene expression, we examined the attention maps. HisToGene’s modified Vision Transformer model has 8 layers and each layer has 16 heads, leading to 128 attention maps. For visualization, we randomly chose the attention map from the first, fourth, and eighth layer, which represent the shallow, median, and deep layers. **Figure 5** shows three representative attention maps when HisToGene predicts the expression for a given target spot (the yellow spot in each plot) under the original spot level resolution and the super-resolution settings. The results indicate that 1) in the shallow layer, the modified Vision Transformer model mainly pays attention to the target spot; 2) in the median layer, the model starts to pay attention to some distant spots, and the pattern is especially clear in the super-resolution setting; 3) in the deep layer, the model pays more attention to distant spots that are tumor related. During the model training process, HisToGene can adaptively change the scale of weights when the input sample size changes; for example, the average weight of the super-resolution attention is about 1/10 of the original-resolution attention. It is also evident that in the super-resolution setting, the model utilizes more refined information provided by the neighboring patches. The difference in the attention weights for input with different sample sizes explains why the gene expression prediction for the same image patch can be different when the number of patches is different.

**Figure 5.**
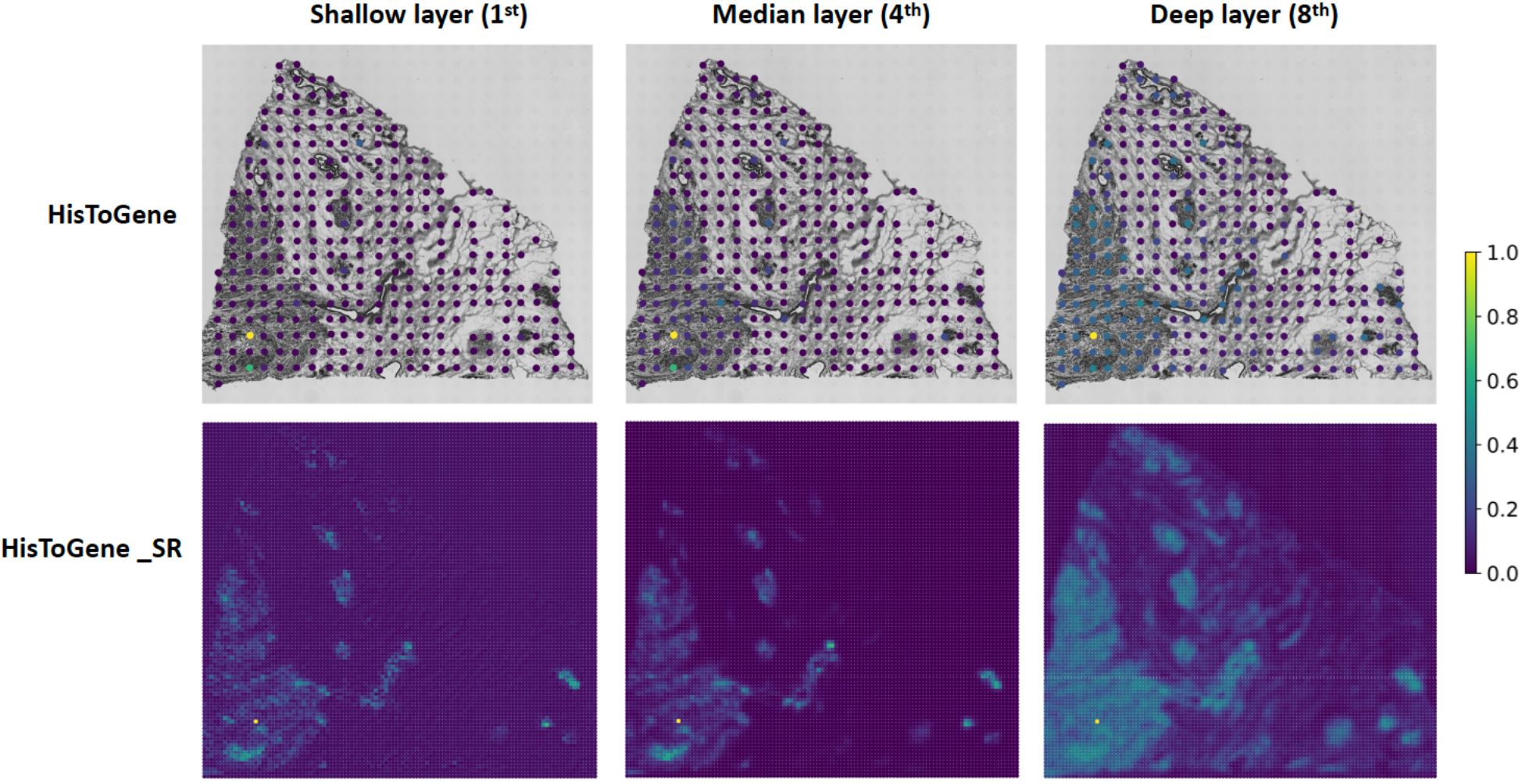
Attention maps in HisToGene’s modified Vision Transformer. Displayed are the attention weights in the modified Vision Transformer when making gene expression predictions for the target spot (the yellow spot in each figure) in the HER2+ breast cancer dataset. The first row shows the attention maps when predicting the gene expression at the original patch/spot level. The second row shows the attention maps when predicting the gene expression at the super-resolution level.

## DISCUSSION

In this paper, we presented HisToGene, a deep learning method that predicts super-resolution gene expression from histology images in tumors. Trained in a ST dataset, HisToGene models the spatial dependency in gene expression and histological features among spots through a modified Vision Transformer model. HisToGene has been evaluated in 32 heterogeneous HER2+ breast cancer tissue sections with 9,612 spots and 785 genes obtained from different patients. Our results consistently show that HisToGene outperformed ST-Net in the spot level gene expression prediction. Additionally, HisToGene can predict gene expression at super-resolution, a feature that ST-Net does not have. To the best of our knowledge, HisToGene is the first method for super-resolution gene expression prediction using histology images. The subsequent clustering analysis using predicted gene expression also shows that HisToGene consistently yielded higher ARIs than ST-Net, and for many of the tissue sections that we analyzed, the ARIs were even higher than those obtained from the observed gene expression. This is likely due to the use of attention, which has the ability to naturally learn from neighborhood. Since the predicted gene expression is based on the histology images, which do not have tissue gaps, it is possible that the consideration of all captured tissue areas in the prediction helped recover expression patterns that are not captured in the observed gene expression.

Compared to ST-Net, HisToGene benefits from the consideration of spots’ dependency and the advanced network architecture, which makes HisToGene robust to heterogeneity among patients. Being robust to batch effects, especially the systematic differences between the training and testing data is an advantage of HisToGene because due to experimental and technical constraints, batch effects are often unavoidable in real studies. HisToGene is robust to heterogeneity among patients due to the following reasons. First, the multi-head attention matrix in HisToGene utilizes the histological features from all spots, implying that when predicting the gene expression for one spot, image features from neighboring spots also contribute. Furthermore, the attention matrix is updated during the training stage, which ensures appropriate adjustment of the neighboring spots’ contributions. Second, HisToGene predicts the gene expression for all spots within a tissue section together. These mechanisms enable HisToGene to model the relationship between histology images and the spatial gene expression data for an entire tissue section, hence minimizing batch effects in histology and gene expression features when learning their relationships. By contrast, CNN-based models such as ST-Net consider each spot independently, making these models more sensitive to batch effects.

HisToGene is computationally fast. To show the computational advantages of HisToGene, we compared its running time for training 31 tissue sections of HER2+ dataset with ST-Net. Our experiment was conducted on CentOS 7 with 24 cores Intel Xeon 8260 CPU and a single NVIDIA V100 (32GB) GPU. On overage, it took HisToGene 11 minutes but 27 minutes for ST-Net.

We mainly focused our analyses on the HER2+ breast cancer dataset in this paper, because this dataset involves a relatively large number of tissue sections and patients. It provides an opportunity to evaluate the performance of HisToGene and ST-Net in the presence of patient heterogeneity. To show the generalizability of HisToGene to other data, we also analyzed a cutaneous squamous cell carcinoma (cSCC) dataset (37), which includes 12 tissue sections obtained from 4 patients, with each patient having 3 sections. Unlike the HER2+ breast cancer dataset, where all tissue sections were generated using the same ST platform, the 12 tissue sections in the cSCC data were generated using a mixture of the relatively low-resolution Spatial Transcriptomics and the more recent 10x Visium platforms. Using the same filtering criteria as the HER2+ breast cancer dataset, 6,630 spots and 134 genes remained for model training and prediction in the cSCC dataset. We also conducted the leave-one-out cross-validation experiment in this dataset, and the results are shown in **Supplementary Note 1**. Due to the relatively small number of tissue sections and patients and the platform heterogeneity among the 12 tissue sections, neither HisToGene nor ST-Net can reliably predict the gene expression with high accuracy. However, HisToGene still yielded higher prediction accuracy than ST-Net. While the requirement of a relatively large training set is a potential limitation of deep learning-based models such as HisToGene, we anticipate that as more and more training ST data become available in the near future, the performance and robustness of HisToGene can be further improved.

## ACKNOWLEDGEMENTS

The authors would like to thank Alma Andersson and Joakim Lundberg for providing the histology images for the HER2+ breast cancer dataset.

## AUTHOR CONTRIBUTIONS

This study was conceived of and led by M.L.. M.P. designed the model and algorithm. M.P. implemented the HisToGene software and led the data analysis with input from M.L. and K.S.. M.P., K.S., and M.L. wrote the paper.

## DATA AVAILABILITY

We analyzed two publicly available ST datasets. These data were acquired from the following websites or accession numbers: (1) human HER2-positive breast tumor ST data (https://github.com/almaan/her2st); (2) human cutaneous squamous cell carcinoma 10x Visium data (GSE144240).

## SOFTWARE AVAILABILITY

An open-source implementation of HisToGene can be downloaded from https://github.com/maxpmx/HisToGene

