## Supplementary Information for "Leveraging information in spatial transcriptomics to predict super-resolution gene expression from histology images in tumors"

**Supplementary Table 1. Top 50 predicted genes by HisToGene.** The genes were ranked by the average -log10 p-values across the 32 tissue sections in the HER2+ breast cancer dataset. For each tissue section, the p-value was obtained by testing whether the correlation between the predicted and observed gene expression was significantly different from zero.

| Rank | Gene | Average -log10 p-values | Average R | Average p-values |
| --- | --- | --- | --- | --- |
| 1 | *GNAS* | 9.33170122 | 0.31840829 | 0.05237465 |
| 2 | *FASN* | 7.33938454 | 0.27453748 | 0.077203 |
| 3 | *MYL12B* | 7.79595066 | 0.27432449 | 0.06200726 |
| 4 | *SCD* | 7.00331428 | 0.26960089 | 0.08393486 |
| 5 | *CLDN4* | 7.03801868 | 0.26145722 | 0.08084089 |
| 6 | *RHOB* | 6.34678558 | 0.23550877 | 0.08655021 |
| 7 | *CCT4* | 6.29548453 | 0.22630695 | 0.09858922 |
| 8 | *TXNDC17* | 5.63395972 | 0.22608913 | 0.10256494 |
| 9 | *SRSF1* | 5.83861687 | 0.2249956 | 0.07870347 |
| 10 | *NDUFB2* | 5.6362317 | 0.22111305 | 0.1297825 |
| 11 | *STMN1* | 4.89820153 | 0.21941921 | 0.08992688 |
| 12 | *ARF6* | 4.75706677 | 0.21931304 | 0.07991021 |
| 13 | *FN1* | 6.20787577 | 0.21876488 | 0.1382412 |
| 14 | *TMEM123* | 6.12693443 | 0.21840646 | 0.0689071 |
| 15 | *ITGB6* | 5.33573753 | 0.21611775 | 0.10550078 |
| 16 | *MID1IP1* | 5.34589362 | 0.21293997 | 0.11831545 |
| 17 | *CRACR2B* | 4.88489549 | 0.21220853 | 0.12390668 |
| 18 | *PDCD5* | 4.5731087 | 0.20737644 | 0.10976367 |
| 19 | *HNRNPUL2* | 4.68590287 | 0.20702539 | 0.14810192 |
| 20 | *TMBIM6* | 4.47153912 | 0.20698966 | 0.08467216 |
| 21 | *CPNE1* | 4.81428105 | 0.20585398 | 0.1004875 |
| 22 | *TMEM14B* | 4.83695732 | 0.20394747 | 0.1544253 |
| 23 | *SRRT* | 4.07371434 | 0.20055954 | 0.08176129 |
| 24 | *SRSF5* | 4.71500809 | 0.19812533 | 0.10558437 |
| 25 | *MAGEF1* | 3.96254834 | 0.19380275 | 0.07907682 |
| 26 | *TNC* | 4.45360785 | 0.19335289 | 0.14747054 |
| 27 | *FADS2* | 4.15906037 | 0.19150386 | 0.11302123 |
| 28 | *UBAP2L* | 4.52071718 | 0.19077351 | 0.15595745 |
| 29 | *FAM193B* | 4.61700395 | 0.18995718 | 0.12005493 |
| 30 | *HMGB2* | 4.40873002 | 0.189363 | 0.12626662 |
| 31 | *GNL2* | 3.76998842 | 0.18862268 | 0.07381618 |
| 32 | *INTS8* | 3.90489731 | 0.18850064 | 0.08032781 |
| 33 | *PRKCSH* | 4.41852901 | 0.18749844 | 0.12276225 |
| 34 | *UFD1L* | 3.78636204 | 0.18711763 | 0.11867428 |
| 35 | *MRPL51* | 4.34005555 | 0.1837053 | 0.19783908 |
| 36 | *NDUFA1* | 4.51502713 | 0.1802292 | 0.1743994 |
| 37 | *NDUFB3* | 4.15071266 | 0.17977559 | 0.13037611 |
| 38 | *KMT2B* | 3.75823539 | 0.17942085 | 0.17293327 |
| 39 | *FOXP4* | 3.62405436 | 0.17714313 | 0.15635171 |
| 40 | *SNRPD3* | 3.54108778 | 0.17614467 | 0.09432443 |
| 41 | *MRPL21* | 3.61855162 | 0.17593202 | 0.14439472 |
| 42 | *MARS* | 3.97667439 | 0.17563173 | 0.14139262 |
| 43 | *LUC7L3* | 3.5480503 | 0.17561638 | 0.12221557 |
| 44 | *DNAJC1* | 3.93216204 | 0.17539909 | 0.20588227 |
| 45 | *CLDN3* | 3.65239902 | 0.17535921 | 0.12612634 |
| 46 | *ATP6AP1* | 4.24316221 | 0.17510715 | 0.2055264 |
| 47 | *SEMA4B* | 3.580602 | 0.17422761 | 0.10177206 |
| 48 | *ATP5O* | 4.22764515 | 0.17363375 | 0.16554514 |
| 49 | *NDUFC1* | 3.49363524 | 0.17213479 | 0.11906831 |
| 50 | *KDM4B* | 3.20692895 | 0.1702299 | 0.15341721 |

**Supplementary Table 2. Top 50 predicted genes by HisToGene*.** The genes were ranked by the average -log10 p-values across the 32 tissue sections in the HER2+ breast cancer dataset. For each tissue section, the p-value was obtained by testing whether the correlation between the predicted and observed gene expression was significantly different from zero.

| Rank | Gene | Average -log10 p-values | Average R | Average p-values |
| --- | --- | --- | --- | --- |
| 1 | *GNAS* | 7.61977783 | 0.26870379 | 0.06576641 |
| 2 | *MYL12B* | 7.06268216 | 0.24256377 | 0.09338287 |
| 3 | *FN1* | 7.32538462 | 0.23929272 | 0.09645939 |
| 4 | *FASN* | 6.36806512 | 0.23626728 | 0.0630492 |
| 5 | *SCD* | 6.00583485 | 0.23044383 | 0.12707601 |
| 6 | *STMN1* | 6.21367867 | 0.22667853 | 0.10912121 |
| 7 | *CLDN4* | 5.97736334 | 0.22604453 | 0.10124676 |
| 8 | *TMEM123* | 6.31249314 | 0.22337847 | 0.11666281 |
| 9 | *RHOB* | 6.29227017 | 0.22060461 | 0.11025758 |
| 10 | *CCT4* | 6.24780454 | 0.21788697 | 0.15961743 |
| 11 | *TXNDC17* | 6.14144215 | 0.21650475 | 0.14453201 |
| 12 | *HMGB2* | 5.93218474 | 0.21542675 | 0.15848342 |
| 13 | *NDUFB2* | 6.01305625 | 0.21188955 | 0.0795658 |
| 14 | *SRSF1* | 5.16403048 | 0.20096756 | 0.08254611 |
| 15 | *MID1IP1* | 5.31748952 | 0.20091082 | 0.15372368 |
| 16 | *PDCD5* | 4.94846744 | 0.19944099 | 0.12544403 |
| 17 | *CPNE1* | 4.88914035 | 0.19402884 | 0.16865547 |
| 18 | *NDUFB3* | 4.92552265 | 0.19205781 | 0.13291532 |
| 19 | *ARF6* | 4.39509704 | 0.19159104 | 0.08635273 |
| 20 | *TMEM14B* | 5.02553197 | 0.19036812 | 0.16784323 |
| 21 | *TEX2* | 4.70471263 | 0.18958416 | 0.14217544 |
| 22 | *HNRNPUL2* | 4.61495301 | 0.18954021 | 0.12622356 |
| 23 | *MRPL51* | 4.74129091 | 0.18851589 | 0.18302173 |
| 24 | *TNC* | 4.77243716 | 0.18845986 | 0.18717087 |
| 25 | *TMBIM6* | 4.64432795 | 0.18786041 | 0.17555732 |
| 26 | *MAGEF1* | 3.65619283 | 0.18507689 | 0.10764109 |
| 27 | *GPRC5A* | 4.64709837 | 0.18506428 | 0.14885693 |
| 28 | *ITGB6* | 4.28398666 | 0.18416358 | 0.10707242 |
| 29 | *NDRG1* | 4.88814068 | 0.18412426 | 0.20451712 |
| 30 | *UBAP2L* | 4.38620805 | 0.18033516 | 0.13326003 |
| 31 | *PRKCSH* | 4.17237876 | 0.18017368 | 0.16260448 |
| 32 | *TCEA3* | 4.29158844 | 0.17927089 | 0.16059555 |
| 33 | *UFD1L* | 4.02791206 | 0.17679789 | 0.17644663 |
| 34 | *CRACR2B* | 4.11746942 | 0.17548713 | 0.16478214 |
| 35 | *NDUFA12* | 3.94143151 | 0.1747079 | 0.17211069 |
| 36 | *GANAB* | 3.94900186 | 0.17308388 | 0.19650422 |
| 37 | *FADS2* | 3.65646226 | 0.17117686 | 0.13705767 |
| 38 | *CRABP2* | 4.33640074 | 0.16989998 | 0.16301158 |
| 39 | *KDM4B* | 3.64161588 | 0.16986658 | 0.17093832 |
| 40 | *BTG1* | 4.04376521 | 0.16903766 | 0.16325509 |
| 41 | *PLXNA1* | 3.54470262 | 0.16885502 | 0.12891258 |
| 42 | *SRRT* | 3.40093416 | 0.16833884 | 0.15881344 |
| 43 | *MARS* | 4.09965713 | 0.1681575 | 0.14276513 |
| 44 | *KMT2B* | 3.5805942 | 0.16809287 | 0.19177506 |
| 45 | *SNRPD3* | 3.99375046 | 0.16740781 | 0.07327653 |
| 46 | *FAM193B* | 3.80964635 | 0.1672208 | 0.11975374 |
| 47 | *GATA3* | 4.08691318 | 0.16714379 | 0.22564067 |
| 48 | *EZH2* | 3.53020505 | 0.16604299 | 0.17369514 |
| 49 | *ATP5H* | 3.56176099 | 0.16598291 | 0.13914845 |
| 50 | *ZFAND3* | 3.40727373 | 0.16565979 | 0.11855663 |

**Supplementary Table 3. Top 50 predicted genes by ST-Net.** The genes were ranked by the average -log10 p-values across the 32 tissue sections in the HER2+ breast cancer dataset. For each tissue section, the p-value was obtained by testing whether the correlation between the predicted and observed gene expression was significantly different from zero.

| Rank | Gene | Average -log10 p-values | Average R | Average p-values |
| --- | --- | --- | --- | --- |
| 1 | *IGHM* | 2.82726396 | 0.07972932 | 0.20749497 |
| 2 | *PTOV1* | 0.84919157 | 0.05915395 | 0.34854101 |
| 3 | *TRAP1* | 0.98583446 | 0.05581523 | 0.29183242 |
| 4 | *DCAF6* | 0.56473981 | 0.04881092 | 0.37765596 |
| 5 | *ATP5O* | 0.90862194 | 0.04811211 | 0.24445701 |
| 6 | *MYL12B* | 0.73968911 | 0.04570592 | 0.33268825 |
| 7 | *SLC35A2* | 0.62459834 | 0.04547275 | 0.33985595 |
| 8 | *NDUFB2* | 0.93111914 | 0.04474541 | 0.2619681 |
| 9 | *SCD* | 0.66578077 | 0.04471691 | 0.36782715 |
| 10 | *MAGOHB* | 0.58429871 | 0.04464906 | 0.31677996 |
| 11 | *PRKCSH* | 0.93898082 | 0.04403122 | 0.28974855 |
| 12 | *TESMIN* | 0.5858354 | 0.04402081 | 0.39624377 |
| 13 | *FAH* | 0.78941295 | 0.04269161 | 0.28449487 |
| 14 | *CLDN4* | 0.78474011 | 0.04255386 | 0.30465686 |
| 15 | *COX5A* | 0.70231815 | 0.04254649 | 0.2989647 |
| 16 | *FOXP4* | 0.69370085 | 0.04156338 | 0.30736544 |
| 17 | *UBAP2L* | 0.61123886 | 0.04075275 | 0.35451713 |
| 18 | *LAMTOR2* | 0.69076053 | 0.04027795 | 0.35372901 |
| 19 | *PAQR4* | 0.57460719 | 0.04023549 | 0.36044515 |
| 20 | *ERAL1* | 0.62380935 | 0.04015197 | 0.30393994 |
| 21 | *HAGHL* | 0.59894346 | 0.03961982 | 0.40428354 |
| 22 | *ENPP1* | 0.49591703 | 0.0393893 | 0.37393832 |
| 23 | *IGHG3* | 0.9575497 | 0.03905437 | 0.32569221 |
| 24 | *DLG1* | 0.56767008 | 0.03882923 | 0.32627679 |
| 25 | *CGGBP1* | 0.67104762 | 0.03854894 | 0.38538718 |
| 26 | *IGLC7* | 0.99113212 | 0.03820342 | 0.27942492 |
| 27 | *TMEM14B* | 0.5808435 | 0.03819166 | 0.41942647 |
| 28 | *HLA-DRA* | 0.94590226 | 0.03797083 | 0.34772444 |
| 29 | *GSK3A* | 0.66270536 | 0.0376525 | 0.41273917 |
| 30 | *TXNRD2* | 0.53442252 | 0.03741065 | 0.35509564 |
| 31 | *PRPF6* | 0.64526157 | 0.03698679 | 0.37562551 |
| 32 | *DNAJB2* | 0.56650873 | 0.03682871 | 0.38014321 |
| 33 | *LTF* | 0.77046644 | 0.03617348 | 0.41850936 |
| 34 | *MUC1* | 0.97278273 | 0.03615715 | 0.33548772 |
| 35 | *GNL2* | 0.60189108 | 0.03612749 | 0.45632277 |
| 36 | *ETFB* | 0.94596691 | 0.03603736 | 0.34347255 |
| 37 | *PNMT* | 1.78910807 | 0.0359639 | 0.33250959 |
| 38 | *SIGMAR1* | 0.52396396 | 0.03582182 | 0.34878753 |
| 39 | *PNPLA6* | 0.59401502 | 0.03579971 | 0.37565066 |
| 40 | *GBP1* | 0.55737405 | 0.03573509 | 0.42039842 |
| 41 | *MST1R* | 0.63480167 | 0.03542165 | 0.3184916 |
| 42 | *QSOX1* | 0.77540602 | 0.03502413 | 0.35112316 |
| 43 | *CRACR2B* | 0.6170485 | 0.03494882 | 0.41456623 |
| 44 | *RNF135* | 0.6696558 | 0.03492154 | 0.31704758 |
| 45 | *C19orf52* | 0.58441118 | 0.03476694 | 0.40497835 |
| 46 | *TCEA2* | 0.5231705 | 0.03444116 | 0.3873315 |
| 47 | *TMEM123* | 0.76339184 | 0.03374817 | 0.33128481 |
| 48 | *COL4A1* | 0.7660625 | 0.03346372 | 0.35964768 |
| 49 | *PER1* | 0.48522731 | 0.03320229 | 0.42741954 |
| 50 | *NDUFS1* | 0.55562443 | 0.03299725 | 0.37496126 |

**Supplementary Figure 1: Visualization of the top 4 genes predicted by ST-Net in the HER2+ breast cancer dataset.** The genes were selected based on the average -log10 p-values across all 32 tissue sections, where the p-value for each tissue section was obtained by testing whether the correlation between the predicted and observed gene expression was significantly different from zero. For each of the 4 genes, the tissue section that had the smallest p-value by ST-Net was selected for visualization. HisToGene* was based on the recovered patch/spot-level gene expression obtained from the super-resolution gene expression prediction, denoted by HisToGene_SR.

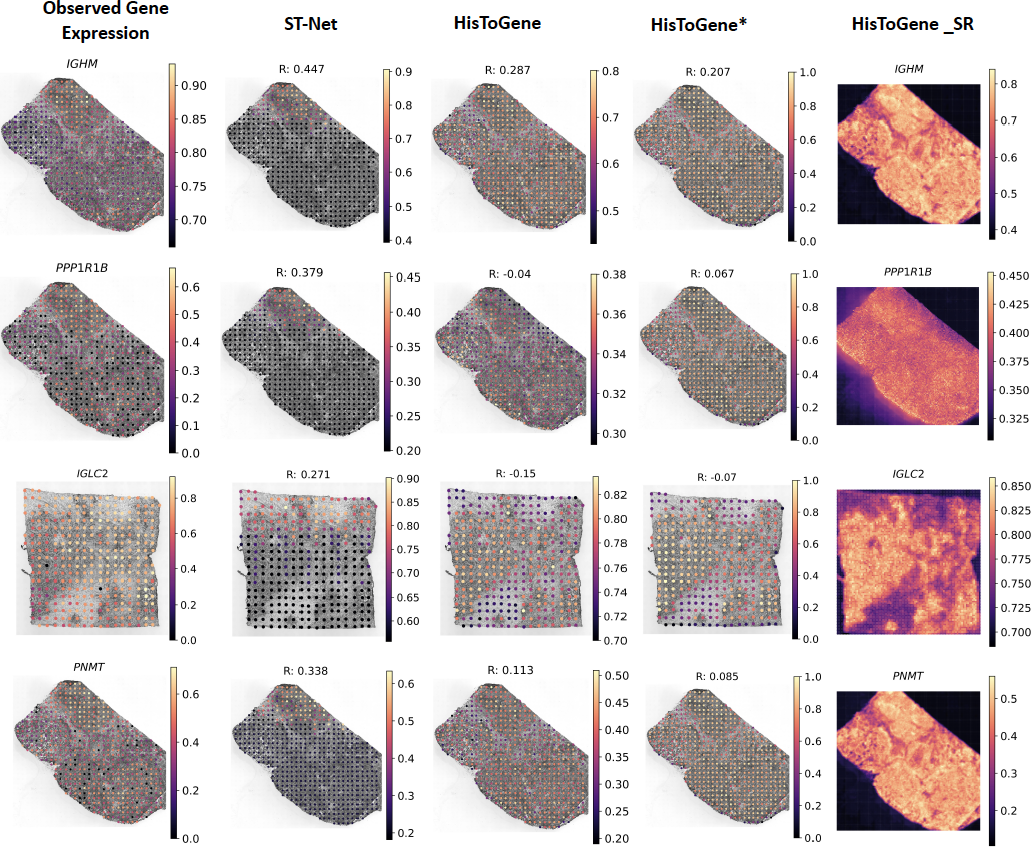

**Supplementary Note 1: Analysis of the cutaneous squamous cell carcinoma (cSCC) dataset.**

This dataset was generated by Ji et al. (1). It includes 12 tissue sections obtained from 4 patients, with each patient having 3 sections. Using the same filtering method, 134 genes remained for expression prediction in the cSCC dataset. We also conducted the leave-one-out cross-validation experiment in this dataset. Results from our analyses are shown in **Supplementary Figures 2-5.**

**Supplementary Figure 2:** Boxplot of the Pearson correlations between the predicted and observed gene expression for the 134 genes predicted by HisToGene, HisToGene*, and ST-Net. HisToGene* is based on the recovered patch/spot-level gene expression obtained from the super-resolution gene expression prediction, denoted by HisToGene_SR.

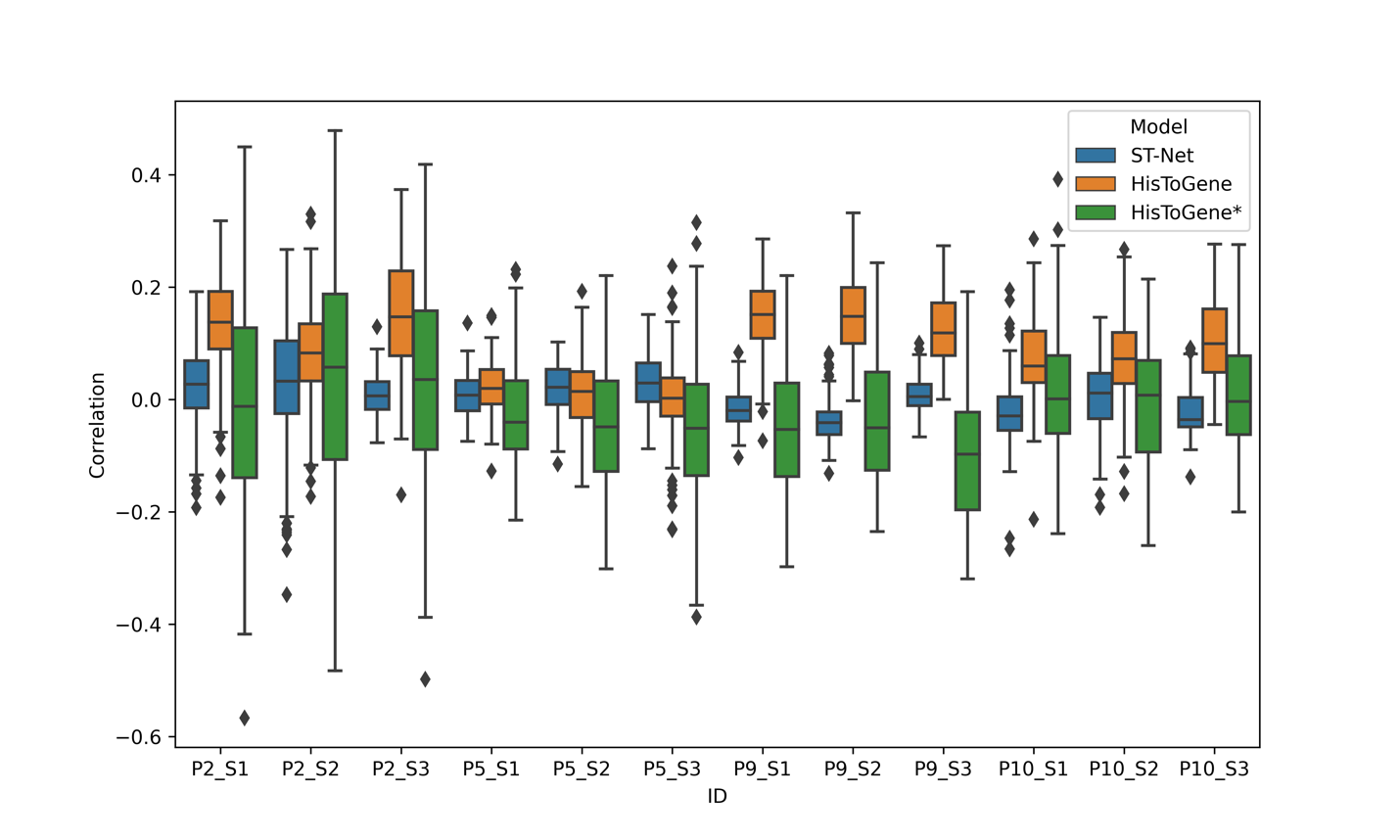

**Supplementary Fig. 3:** **Visualization of the top 4 genes predicted by HisToGene in the cSCC dataset.** The genes were selected based on the average -log10 p-values across all 12 tissue sections, where the p-value for each tissue section was obtained by testing whether the correlation between the predicted and observed gene expression was significantly different from zero. For each of the 4 genes, the tissue section that had the smallest p-value by ST-Net was selected for visualization. HisToGene* was based on the recovered patch/spot-level gene expression obtained from the super-resolution gene expression prediction, denoted by HisToGene_SR.

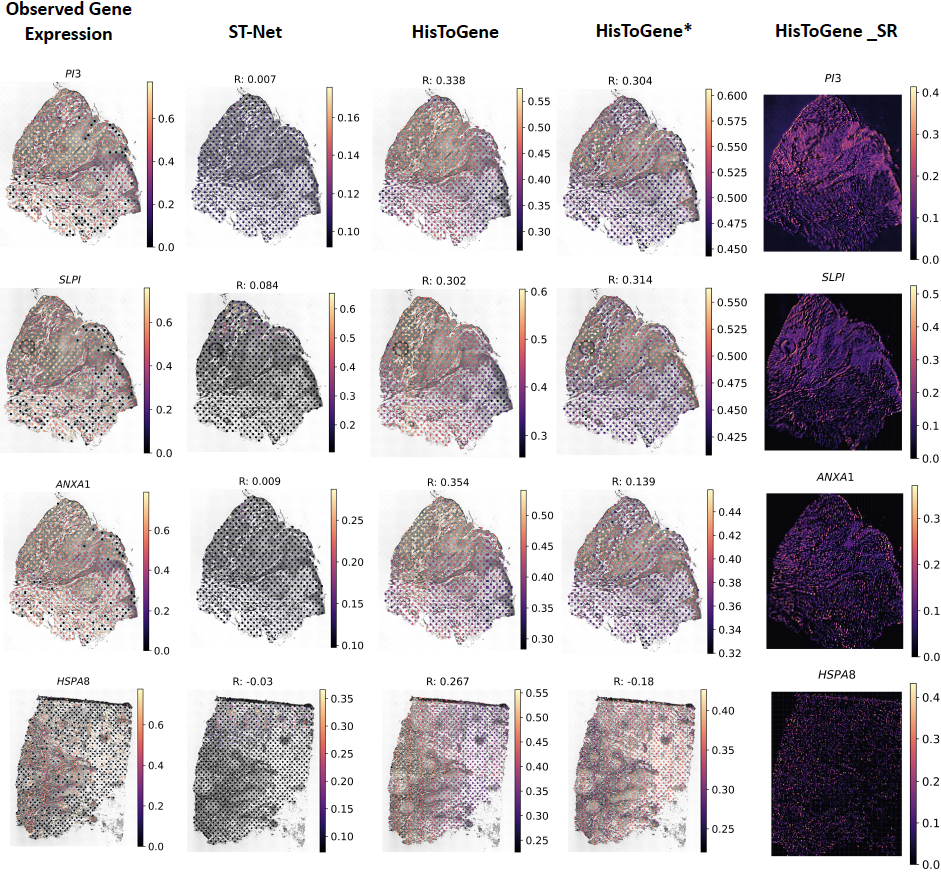

**Supplementary Fig. 4: Visualization of the top 4 predicted genes by HisToGene* in the cSCC dataset.** The genes were selected based on the average -log10 p-values across all 12 tissue sections, where the p-value for each tissue section was obtained by testing whether the correlation between the predicted and observed gene expression was significantly different from zero. For each of the 4 genes, the tissue section that had the smallest p-value by ST-Net was selected. HisToGene* was based on the recovered patch/spot-level gene expression obtained from the super-resolution gene expression prediction, denoted by HisToGene_SR.

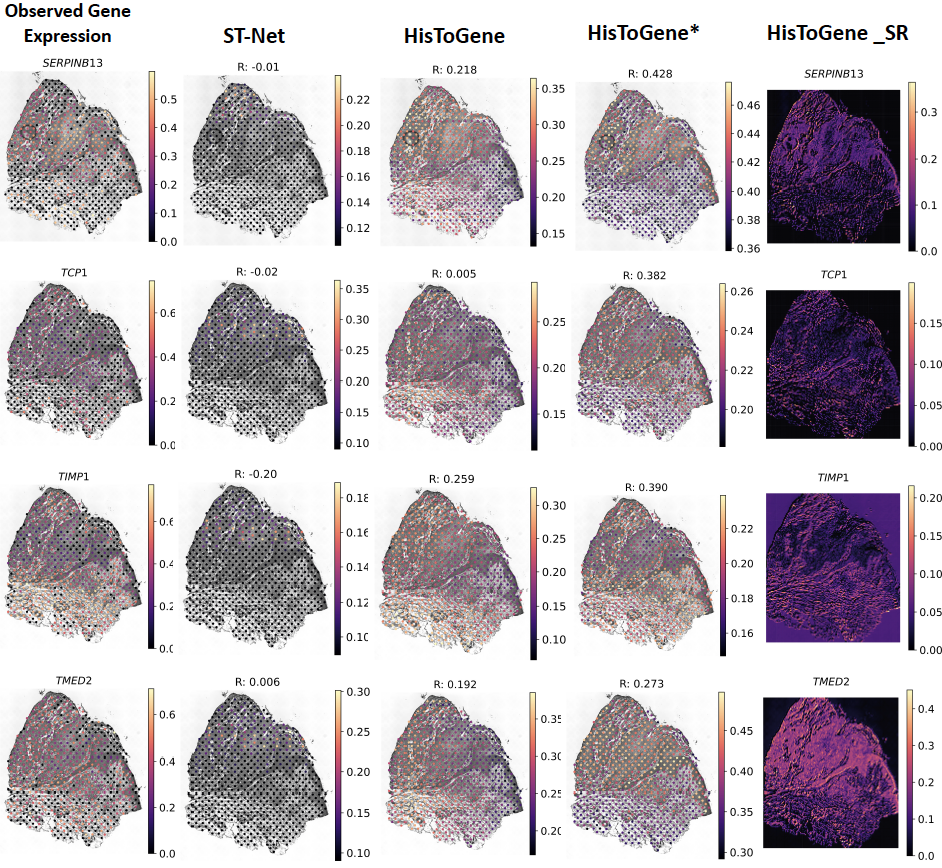

**Supplementary Fig. 5: Visualization of the top 4 predicted genes by ST-Net in the cSCC dataset.** The genes were selected based on the average -log10 p-values across all 12 tissue sections, where the p-value for each tissue section was obtained by testing whether the correlation between the predicted and observed gene expression was significantly different from zero. For each of the 4 genes, the tissue section that had the smallest p-value by ST-Net was selected. HisToGene* was based on the recovered patch/spot-level gene expression obtained from the super-resolution gene expression prediction, denoted by HisToGene_SR.

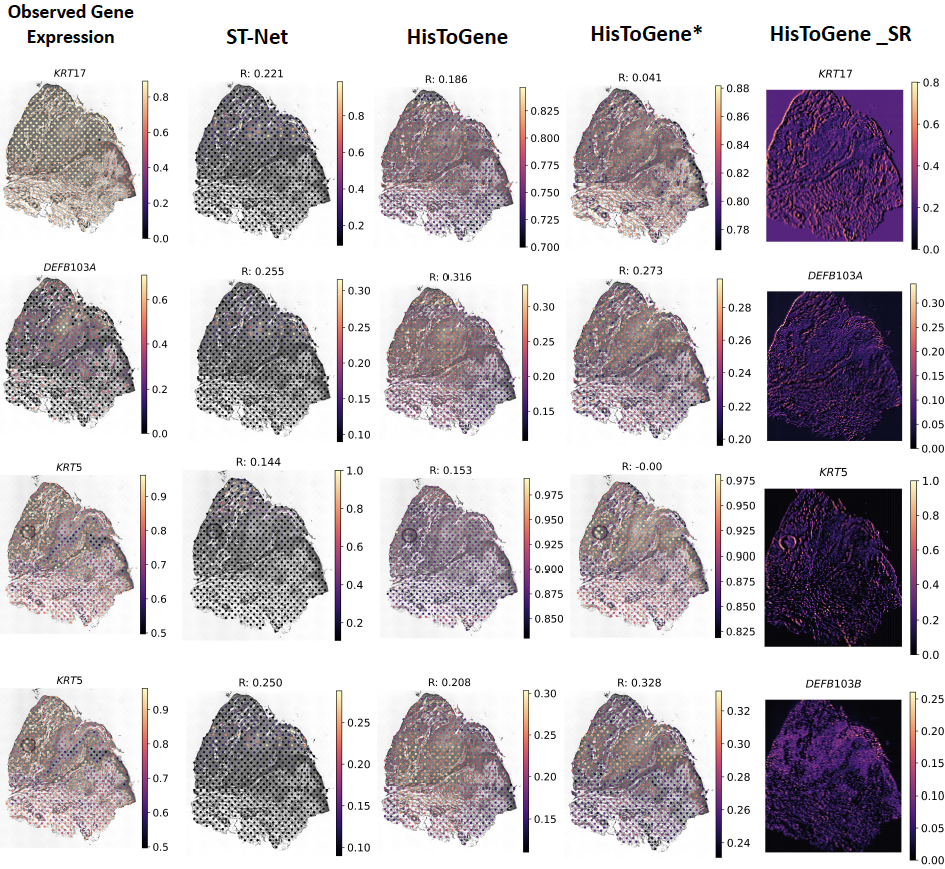
